# Statistical properties of the MetaCore network of protein-protein interactions

**DOI:** 10.1101/2021.04.02.438245

**Authors:** Ekaterina Kotelnikova, Klaus M. Frahm, José Lages, Dima L Shepelyansky

## Abstract

The MetaCore commercial database describes interactions of proteins and other chemical molecules and clusters in the form of directed network between these elements, viewed as nodes. The number of nodes goes beyond 40 thousands with almost 300 thousands links between them. The links have essentially bi-functional nature describing either activation or inhibition actions between proteins. We present here the analysis of statistical properties of this complex network applying the methods of the Google matrix, PageRank and CheiRank algorithms broadly used in the frame of the World Wide Web, Wikipedia, the world trade and other directed networks. We specifically describe the Ising PageRank approach which allows to treat the bi-functional type of protein-protein interactions. We also show that the developed reduced Google matrix algorithm allows to obtain an effective network of interactions inside a specific group of selected proteins. This method takes into account not only direct protein-protein interactions but also recover their indirect nontrivial couplings appearing due to summation over all the pathways passing via the global bi-functional network. The developed analysis allows to espablish an average action of each protein being more oriented to activation or inhibition. We argue that the described Google matrix analysis represents an efficient tool for investigation of influence of specific groups of proteins related to specific diseases.

## 1 Introduction

The MetaCore database [1] provides a large size network of Protein-Protein Interactions (PPI). It has been shown to be useful for analysis of specific biological problems (see e.g. [2, 3]) and finds various medical applications. At present, the network has *N* = 40079 nodes with *N_ℓ_* = 292904 links and an average of *n*_*l*_ = *N*_*l*_/*N* ≈ 7.3 links per node. The nodes are composed mainly by proteins but in addition there are also certain molecules and molecular clusters catalyzing the interactions with proteins. This PPI network is directed and non-weighted. Its interesting feature is the bi-functional nature of the links leading to either the activation or the inhibition of one protein by another one. In some cases, the link action is neutral or unknown.

In the present work, we describe the statistical properties and the Google matrix analysis (GMA) of the MetaCore network. The GMA and the related PageRank algorithm has been at the foundation of the Google search engine with important applications to the World Wide Web (WWW) analysis [4, 5]. A variety of GMA applications to directed networks are presented in [6]. The first application of the GMA to PPI network was reported for the SIGNOR PPI network [7] in [8]. However, the size of the SIGNOR network is by a factor ten smaller than the MetaCore one and thus the GMA of SIGNOR network can be considered only as a test bed for more detailed studies of PPI.

An important feature of the PPI networks is the bi-functional character of the directed links representing activation or inhibition actions. Usually, the directed networks have been considered without functionality of links (see e.g. [4, 5, 6]). The Ising-Google matrix analysis (IGMA) [9] extends the GMA for bi-functional links. A test application to the SIGNOR PPI network [7] can be found in [10]. The Ising-Google matrix analysis (IGMA) represents each node by two states ↑ and ↓, like Ising spins up and down. A link is then represented by a 2 × 2-matrix describing the actions of activation or inhibition [9, 10]. By contrast with the case of links without functionality, this description leads to a doubling of the number of nodes *N*_*I*_ = 2*N*. In the present work, we apply the IGMA to the MetaCore network which provide bi-functional interactions between multiple proteins.

In addition, we also use the reduced Google matrix analysis (RGMA), developed in [11, 12], to describe the effective interactions between a subset of *N*_r_ « *N* selected nodes taking account of all the indirect pathways connecting each couple of these *N*_r_ nodes throughout the global PPI network. The efficiency of the RGMA has been demonstrated for large variety of directed networks including Wikipedia and the world trade network (see e.g. [13, 14]). The RGMA adapted to the IGMA for bi-functional links is called hereafter the RIGMA.

The paper is composed as follows: the data sets and the methods are described in the Section 2, the results are presented in the Section 3 and the discussion and the conclusion are given in the Section 4.

## 2 Data sets and methods

### 2.1 Google matrix construction of the MetaCore network

At the first step, we start the construction of the Google matrix *G* of the MetaCore network neglecting the bi-functional character of the links and considering unweighted links. Considering the adjacency matrix *A*, the elements *A*_*ij*_ of which are equal to 1 if node *j* points to node *i* and equal to 0 otherwise, the stochastic matrix *S* of the node-to-node Markov transitions is obtained by normalizing to unity each column of the adjacency matrix *A*. For dangling nodes, the corresponding column is filled with elements with value 1/*N*. The stochastic matrix *S* describes a Markov chain process on the network: a random surfer hops from node to node in accordance with the network structure and hops anywhere on the network if it reaches a dangling node. The elements of the Google matrix *G* takes then the standard form

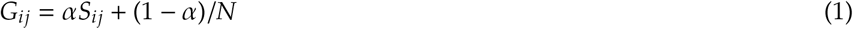

where 0.5 ≤ *α* < 1 is the damping factor. The random surfer obeying to the stochastic process encoded in *G* explores, with a probability *α*, the network in accordance to the stochastic matrix *S* and hops, with a complementary probability (1 − *α*), to any node of the network. The damping factor allows the random surfer to escape from possible isolated communities. Here, we use the standard value *α* = 0.85 [5, 6]. The PageRank vector *P* is the right eigenvector of the Google matrix *G* corresponding to the leading eigenvalue, here *λ* = 1. The corresponding eigenproblem equation is then *GP* = *P*. According to the Perron-Frobenius theorem, the PageRank vector *P* has positive elements. The PageRank vector element *P*(*j*) gives the probability to find the random surfer on the node *j* once the Markov process has reached the stationary regime. Consequently, all the nodes can be ranked by decreasing PageRank probability. We define the PageRank index *K*(*j*) giving the rank of the node *j*. The node *j* with the highest (lowest) PageRank probability *P*(*j*) corresponds to *K*(*j*) = 1 (*K*(*j*) = *N*). On average, the PageRank probability *P*(*j*) is proportional to the number of ingoing links pointing to node *j*.

It is also useful to consider a network obtained by the inversion of all the directions of the links. For this inverted network, the corresponding Google matrix is denoted *G*^*^ and the corresponding PageRank vector is called the CheiRank vector *P*^*^ and is defined such as *G*^*^*P*^*^ = *P*^*^. The importance and the detailed statistical analysis of the CheiRank vector have been reported in [15, 16] (see also [6, 14]). Similarly to the PageRank vector, the CheiRank probability *P*^*^(*j*) is proportional, on average, to the number of outgoing links going out from node *j*. We define also a CheiRank index *K*^*^(*j*) giving the rank of the node *j* according to its CheiRank probability *P*^*^(*j*).

### 2.2 Reduced Google matrix

The concept of the reduced Google matrix analysis (RGMA) was introduced in [11] and applied with details to Wikipedia networks in [12]. The RGMA determines effective interactions between a selected subset of *N*_r_ nodes embedded in a global network of size *N* ≫ *N*_r_. These effective interactions are determined taking into account that there are many indirect links between the *N*_r_ nodes via all the other *N*_s_ = *N* − *N*_r_ nodes of the network. As an example, we may have two nodes *A* and *C* which belongs to the selected subset of *N*_r_ nodes and which are not coupled by any direct link. However, it may exist a chain of links from *A* to *B*_1_, then from *B*_1_ to *B*_2_,…, and then from *B*_*m*_ to *C* where *B*_1_, …, *B*_*m*_ are nodes not belonging to the subset of *N*_r_ nodes. Although *A* and *C* are not directly connected, the is a chain of *m* + 1 directed links indirectly connecting *A* and *C*. The RGMA allows to infer an effective weighted link between any couple of two nodes of the *N*_r_ subset of interest taking account of the possible direct link existing between these two nodes and taking account of all the possible chains of links connecting them throughout the remaining global network of size *N*_s_ = *N* − *N*_r_ ≫ *N*_r_. It is important to stress that rather often the network analysis is done taking only into account the direct links between the *N*_r_ nodes and, as a consequence, completely omitting their indirect interactions via the global network. It is known that such a simplified approach produces erroneous results as it happened for the network of historical figures extracted from Wikipedia when only direct links between historical figures were taking into account and all other links had been omitted [17] (see discussion at [18]).

It is convenient to write the Google matrix *G* associated to the global network as

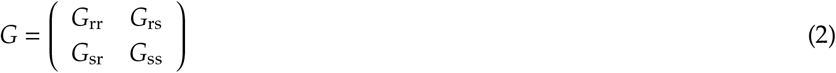

where the label “r” refers to the nodes of the reduced network, ie the subset of *N*_r_ nodes, and “s” to the other *N*_s_ = *N* − *N*_r_ nodes which form the complementary network acting as an effective “scattering network”. The reduced Google matrix *G*_R_ associated to the subset of the *N*_r_ nodes is a *N*_r_ × *N*_r_ matrix defined as

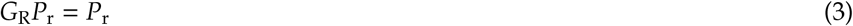

where *P*_r_ is a *N*_r_ size vector the components of which are the normalized PageRank probabilities of the *N*_r_ nodes of interest,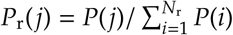. The RGMA consists in finding an effective Google matrix for the subset of *N*_r_ nodes keeping the relative ranking between these nodes. To ensure the relation (3), the reduced Google matrix *G*_R_ has the form [11, 12]

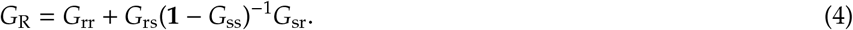

As shown in [11, 12], the reduced Google matrix *G*_R_ can be represented as the sum of three components

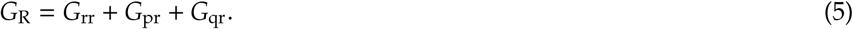

Here, the first component, *G*_rr_, corresponds to the direct transitions between the *N*_r_ nodes; the second component, *G*_pr_, is a matrix of rank with all the columns being approximately equal to the reduced PageRank vector *P*_r_; the third component, *G*_qr_, describes all the indirect pathways passing through the global network. Thus, the component *G*_qr_ represents the most nontrivial information related to indirect hidden transitions. We also define *G*_qrnd_ matrix which is the *G*_qr_ matrix deprived of its diagonal elements. The contribution of each component is characterized by their weights *W*_R_, *W*_pr_, *W*_rr_, *W*_qr_ (*W*_qrnd_) respectively for *G*_R_, *G*_pr_, *G*_rr_, *G*_qr_ (*G*_qrnd_). The weight of a matrix is given by the sum of all the matrix elements divided by its size, here *N*_r_ (by definition *W*_R_ = 1). Examples of reduced Google matrices associated to various directed networks are given in [8, 12, 14, 10].

### 2.3 Bi-functional Ising MetaCore network

To take into account the bi-functional nature (activation and inhibition) of MetaCore links, we use the approach proposed in [9, 10] with the construction of a larger network where each node is split into two new nodes with labels (+) and (−). These two nodes can be viewed as two Ising-spin components associated to the activation and the inhibition of the corresponding protein. To construct the doubled “Ising” network of proteins, each elements of the initial adjacency matrix is replaced by one of the following 2 × 2 matrices

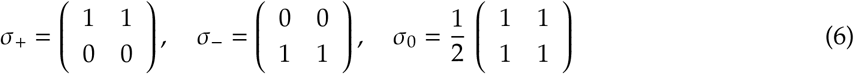

where σ_+_ applies to “activation” links, σ_−_ to “inhibition” links, and σ_0_ when the nature of the interaction is “unknown” or “neutral”. For the rare cases of multiple interactions between two proteins, we use the sum of the corresponding σ-matrices which increases the weight of the adjacency matrix elements. Once the “Ising” adjacency matrix is obtained, the corresponding Google matrix is constructed in the usual way (see Section 2.1). The initial simple MetaCore network has *N* = 40079 nodes and *N_𝓁_* = 292904 links; the ratio of the number of activation/inhibition links is *N*_*𝓁* +_/*N*_*𝓁* −_ = 65379/49384 ≃1.3 and the number of neutral links is *N*_*𝓁n*_ = *N_𝓁_* − *N* _*𝓁*+_ − *N*_*𝓁*−_ = 178141. The doubled Ising MetaCore network corresponds to *N*_*I*_ = 80158 nodes and *N*_*I,𝓁*_ = 942090 links (according to the non-zero entries of the used σ-matrices).

Now, the PageRank vector associated to this doubled Ising network has two components *P*_+_(*j*) and *P*_−_(*j*) for every node *j* of the simple network. Due to the particular structure of the σ-matrices (6), one can show analytically the exact identity, *P*(*j*) = *P*_+_(*j*) + *P*_−_(*j*), where *P*(*j*) is the PageRank of the initial single PPI network. We have numerically verified that the identity *P*(*j*) = *P*_+_(*j*) + *P*_−_(*j*) holds up to the numerical precision ∼ 10^−13^.

As in [9], we characterize each node by its PageRank “magnetization” given by

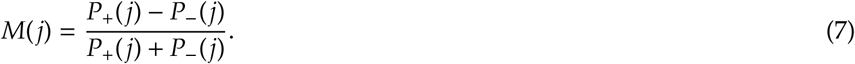

By definition, we have −1 ≤ *M*(*j*) ≤ 1. Nodes with positive *M* are mainly activated nodes and those with negative *M* are mainly inhibited nodes.

### 2.4 Sensitivity

The reduced Google matrix *G*_R_ of bi-functional (or Ising) MetaCore network describes effective interactions between *N*_r_ nodes taking into account the activation or inhibition nature of the interactions.

Following [13], it is useful to determine the sensitivity of the PageRank probabilities in respect to small variation of the matrix elements of *G*_R_. The PageRank sensitivity of the node *j* with respect to a small variation of the *b* → *a* link is defined as

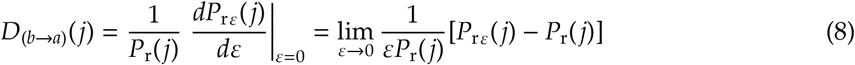

where *P*_rε_(*j*) is the PageRank vector computed from a perturbed matrix *G*_Rε_ the elements of which are defined by *G*_Rε_(*a, b*) = *G*_R_(*a, b*)(1 + ε)/[1 + ε*G*_R_(*a, b*)] for the element (*a, b*), *G*_Rε_(*c, b*) = *G*_R_(*c, b*)/[1 + ε*G*_R_(*a, b*)] for the other elements (*c, b*) in the same column *b*, and *G*_Rε_(*c, d*) = *G*_R_(*c, d*) for the elements (*c, d*) in the other columns. The factor 1/[1 + ε*G*_R_(*a, b*)] ensures the correct sum normalization of the modified column *b*.

We use here an efficient algorithm described in [19] to evaluate the derivative in (8) exactly without usage of finite differences.

As proposed in [13], we define the symmetric matrix (see Eq.15 of [13])

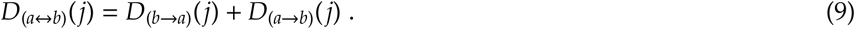

and furthermore the two symmetric and anti-symmetric sensitivity matrices

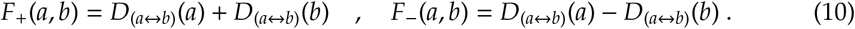

These two sensitivity matrices characterize a variation of PageRank with a small variation of coupling matrix element between *b* and *a* nodes.

## 3 Results

Below, we describe various statistical properties of the MetaCore network obtained by the methods described above. More detailed data are available at [20].

### 3.1 CheiRank and PageRank of the MetaCore network

Let us sort the PageRank probabilities from the highest value to which we associate the *K* = 1 rank to the smallest value to which we associate to the *K* = *N* rank.

The dependence *P*(*K*) of the PageRank probabilities on the PageRank index *K* and the dependence *P*^*^(*K*^*^) of the CheiRank probabilities on the CheiRank index *K*^*^ are shown in Fig. 1 for the simple MetaCore network and the Ising (doubled) MetaCore network. The decay of the probabilities is approximately proportional to an inverse index in a power β ≈ 2/3, ie *P*(*K*) ∝ 1/*K*^2/3^. This exponent β is approximately the same for the PageRank and the CheiRank probabilities, and for both network types. The situation is different from the networks of WWW, Wikipedia, and Linux for which one usually have β ≈ 0.9 for the PageRank probabilities and β ≈ 0.6 for CheiRank probabilities [5, 6, 15]. We assume that in PPI networks both ingoing and outgoing links are of equal importance while, in the other above cited networks, ingoing links are more robust and stable than outgoing links which have a more random character.

**Figure 1.**
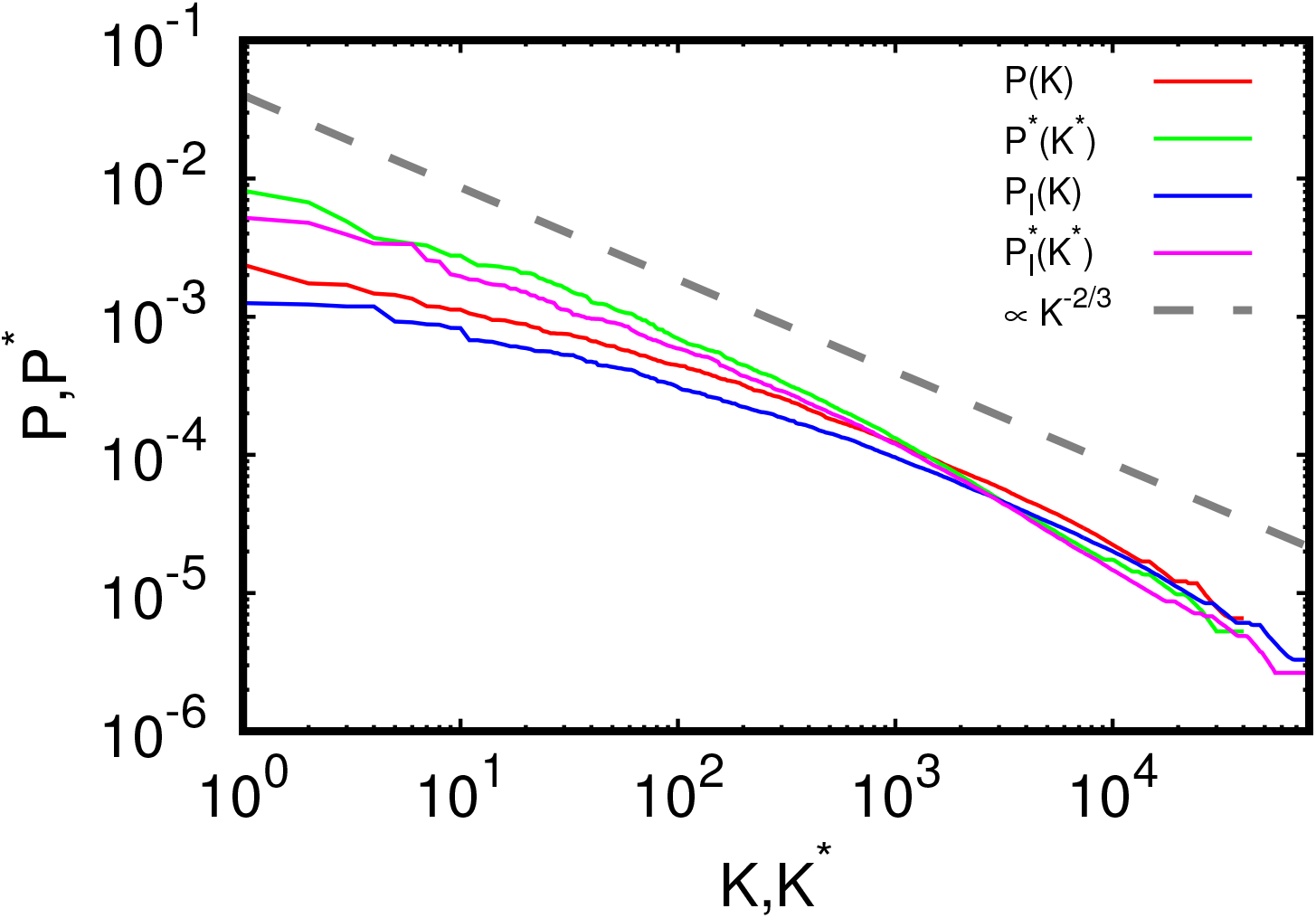
PageRank probability *P*(*K*) (*P*_*I*_(*K*)) and CheiRank probability *P^*^*(*K^*^*) 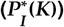 are shown as a function of the corresponding rank indexes *K* and *K*^*^ for the simple (Ising) MetaCore network. For comparison, the dashed gray line corresponds to the power decay *P* ∝ *K*^−2/3^.

The top 40 PageRank and CheiRank nodes of the MetaCore network are given in Tables 1 and 2 respectively. The top 3 PageRank positions are occupied by specific molecules actively participating in various reactions with proteins. The top 3 CheiRank positions are occupied by the transcription factor c-Myc, the generic enzyme eIF2C2 (Argonaute-2), and the generic binding protein IGF2BP3. In a certain sense, we can say that top PageRank nodes are like workers in a company, who receive many orders, while top CheiRank nodes are like company administrators who submit many orders to their workers (such a situation was discussed for a company management network [21]).

**Table 1.**
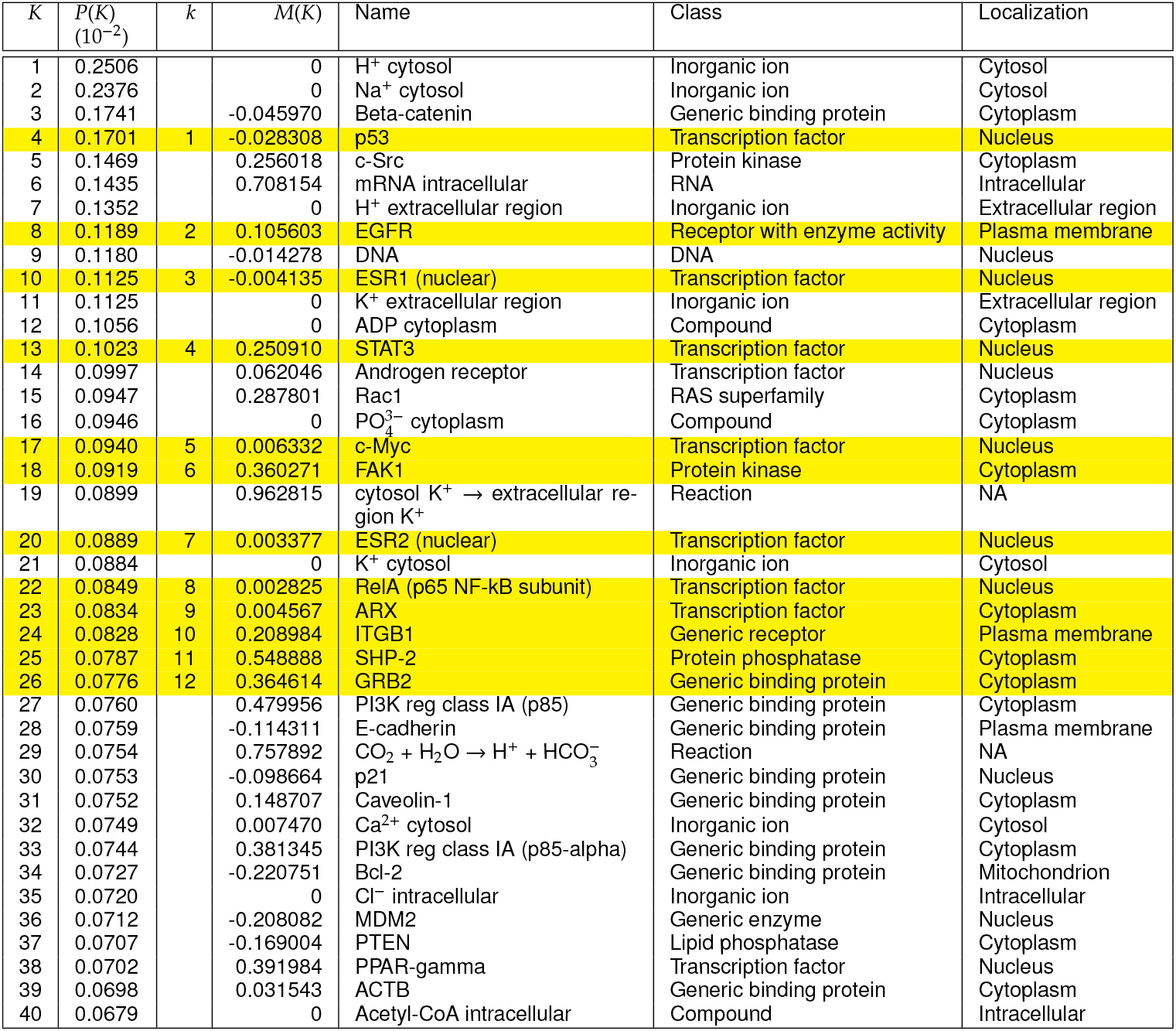
**Top 40 PageRank nodes of the simple MetaCore network. These nodes are sorted by descending PageRank probabilities *P*(*K*) and consequently by ascending PageRank index *K*. The corresponding name, class and bio-localization of the node is given. The values *M*(*K*) of the PageRank magnetization (7) are also given. The node highlighted in yellow corresponds to the twelve proteins selected for the RGMA and RIGMA analysis. These twelve proteins are ordered by the relative PageRank index *k*. Here, NA means not applicable.**

**Table 2.** **Top 40 CheiRank nodes of the simple MetaCore network. These nodes are sorted by descending CheiRank probabilities *P*^*^(*K*^*^) and consequently by ascending PageRank index *K*^*^. The corresponding name, class and bio-localization of the node is given. The values *M*(*K*^*^) of the PageRank magnetization (7) are also given. The node highlighted in green corresponds to proteins from the subset of the twelve proteins chosen in Table 1 with *K*^*^≤ 40. These proteins are ordered by the relative PageRank index *k*^*^.**

| $K^*$ | $P^*(K^*)$<br>($10^{-2}$) | $k^*$ | $M(K^*)$ | Name | Class | Localization |
| --- | --- | --- | --- | --- | --- | --- |
| 1 | 1.1464 | 1 | 0.006332 | c-Myc | Transcription factor | Nucleus |
| 2 | 0.8172 |  | 0.035667 | eIF2C2 (Argonaute-2) | Generic enzyme | Cytoplasm |
| 3 | 0.6722 |  | -0.174071 | IGF2BP3 | Generic binding protein | Cytoplasm |
| 4 | 0.4890 |  | 0.680968 | Ubiquitin | Generic binding protein | Cytoplasm |
| 5 | 0.3719 |  | 0.110759 | SOX9 | Transcription factor | Nucleus |
| 6 | 0.3529 | 2 | -0.028308 | p53 | Transcription factor | Nucleus |
| 7 | 0.3373 |  | 0.228978 | c-Fos | Transcription factor | Nucleus |
| 8 | 0.3276 |  | 0 | CUX1 (p110) | Transcription factor | Nucleus |
| 9 | 0.2989 |  | -0.057557 | SP1 | Transcription factor | Nucleus |
| 10 | 0.2770 | 3 | -0.004135 | ESR1 (nuclear) | Transcription factor | Nucleus |
| 11 | 0.2769 | 4 | 0.002825 | RelA (p65 NF-kB subunit) | Transcription factor | Nucleus |
| 12 | 0.2534 |  | -0.010911 | eIF2C1 (Argonaute-1) | Generic binding protein | Cytoplasm |
| 13 | 0.2354 |  | 0.062046 | Androgen receptor | Transcription factor | Nucleus |
| 14 | 0.2350 |  | -0.045970 | Beta-catenin | Generic binding protein | Cytoplasm |
| 15 | 0.2330 |  | -0.075622 | BRD4 | Generic binding protein | Nucleus |
| 16 | 0.2308 |  | 0.153950 | Oct-3/4 | Transcription factor | Nucleus |
| 17 | 0.2259 |  | -0.001577 | PUM2 | Generic binding protein | Cytoplasm |
| 18 | 0.2239 |  | 0.188479 | EZH2 | Generic enzyme | Nucleus |
| 19 | 0.2193 |  | 0.208146 | p300 | Generic enzyme | Nucleus |
| 20 | 0.2072 |  | -0.407833 | TUG1 | RNA | Cytoplasm |
| 21 | 0.2072 |  | -0.118501 | E2F1 | Transcription factor | Nucleus |
| 22 | 0.2062 |  | 0 | ASCC2 | Generic binding protein | Nucleus |
| 23 | 0.2005 |  | 0 | LIMR | Generic receptor | Plasma membrane |
| 24 | 0.1903 |  | 0.148471 | BRG1 | Generic enzyme | Nucleus |
| 25 | 0.1871 | 5 | 0.250910 | STAT3 | Transcription factor | Nucleus |
| 26 | 0.1811 |  | 0.381258 | RBM24 | Generic binding protein | Cytoplasm |
| 27 | 0.1789 |  | 0.746981 | SUMO-1 | Generic binding protein | Nucleus |
| 28 | 0.1728 |  | 0.140357 | c-IAP2 | Generic binding protein | Cytoplasm |
| 29 | 0.1699 |  | 0.038221 | HIF1A | Transcription factor | Nucleus |
| 30 | 0.1677 |  | 0 | Zn<sup>2+</sup> cytosol | Inorganic ion | Cytosol |
| 31 | 0.1623 |  | -0.013644 | CDK9 | Protein kinase | Cytoplasm |
| 32 | 0.1587 |  | -0.223816 | MeCP2 | Generic binding protein | Nucleus |
| 33 | 0.1533 |  | -0.053592 | ELAVL1 (HuR) | Generic binding protein | Nucleus |
| 34 | 0.1497 |  | 0.120649 | HDAC1 | Generic enzyme | Nucleus |
| 35 | 0.1473 |  | -0.034082 | BRD7 | Generic binding protein | Nucleus |
| 36 | 0.1452 |  | 0.131956 | CREB1 | Transcription factor | Nucleus |
| 37 | 0.1449 |  | 0 | Zn<sup>2+</sup> nucleus | Inorganic ion | Nucleus |
| 38 | 0.1423 |  | 0.096830 | SUMO-2 | Generic binding protein | Cytoplasm |
| 39 | 0.1400 |  | -0.051730 | BRD2 | Protein kinase | Cytoplasm |
| 40 | 0.1343 |  | 0.228824 | C/EBPbeta | Transcription factor | Nucleus |

The density distribution of nodes of the MetaCore network on the PageRank-CheiRank (*K, K*^*^)-plane is shown in Fig. 2. Comparing to the case of Wikipedia networks [6, 16] the distribution is globally more symmetric in respect to the diagonal *K* = *K*^*^. This reflects the fact that the decay of the PageRank and the CheiRank probabilities in Fig. 1 is approximately the same. However, the top nodes are rather different for the PageRank and CheiRank rankings that is also visible from Tables 1 and 2. As an example, the top 40 PageRank and the top 40 CheiRank share only 7 nodes in common (Beta-catenin, p53, ESR1, STAT3, Androgen receptor, c-Myc, RelA) which are transcription factors with the exception of Beta-catenin which is a generic binding protein. As a consequence, depending on the considered biological process, these biological elements trigger the multiple cascades of interactions or are at the very end of these cascades. In contrast, there are biological elements with low *K* and high *K*^*^ and *vice versa*. For example, the phosphate compound 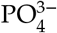, with PageRank-CheiRank indexes (*K* = 16,*K*^*^ = 14888), is mainly a residue of biological processes and the passage of the Potassium ion K+ from the cytosol to the extracellular region, with PageRank-CheiRank indexes (*K* = 19, *K*^*^ = 26346), can be considered as the final step of some biological process.

**Figure 2.**
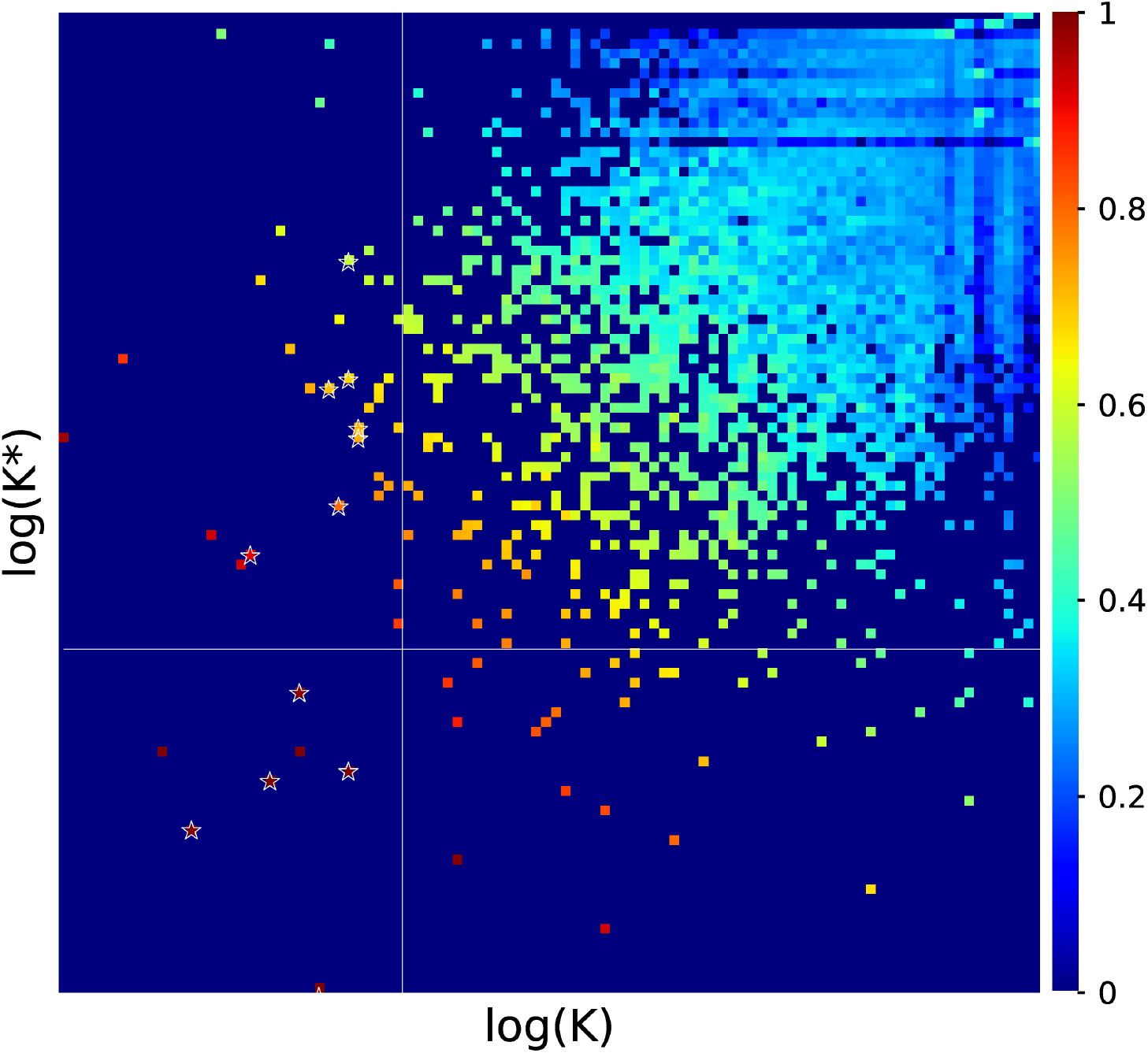
Density of nodes of the MetaCore network on the PageRank-CheiRank (*K, K*^*^)-plane. The numbers of the color bar are a linear function of the logarithm of the density (with maximum values corresponding to 1 (red); minimum non-zero and zero values of the density corresponding to 0 (blue); the distribution is computed for 100 × 100 cells equidistant in logarithmic scale). The white stars indicate the positions of the 12 selected nodes presented in Table 1. The white vertical and horizontal lines represent nodes with *K* ≤ 40 and *K*^*^ ≤ 40, respectively.

Among the top 40 PageRank nodes, we select a subset of 12 nodes which are more directly related to proteins. These 12 nodes are represented by white stars in the Fig. 2. The list of these nodes is given in Table 1. Below, we present the RIGMA analysis of these 12 nodes taking into account of the bi-functionality of the links (activation - inhibition).

### 3.2 Magnetization of nodes of the Ising MetaCore network

From the PageRank probabilities of the Ising MetaCore network, we determine the magnetization *M*(*K*) of each node given by (7). The dependence of the magnetization *M*(*K*) on the PageRank index *K* is shown in Fig. 3. For *K* ≤ 10, only few nodes have a significant positive magnetization. In the range 10 < *K* ≤ 10^3^, some nodes have almost the maximal positive or negative values of the magnetization with *M* being close to 1 or −1. Such nodes perform mainly activation or inhibition actions, respectively. For the range *K* > 10^3^, we see an envelope restricting the maximal or the minimal values *M*. At present, we have no analytical description of this envelope. We suppose that nodes with high *K* values have a majority of outgoing links which are more fluctuating in this range thus giving a decrease of the maximal/minimal values of *M*.

**Figure 3.**
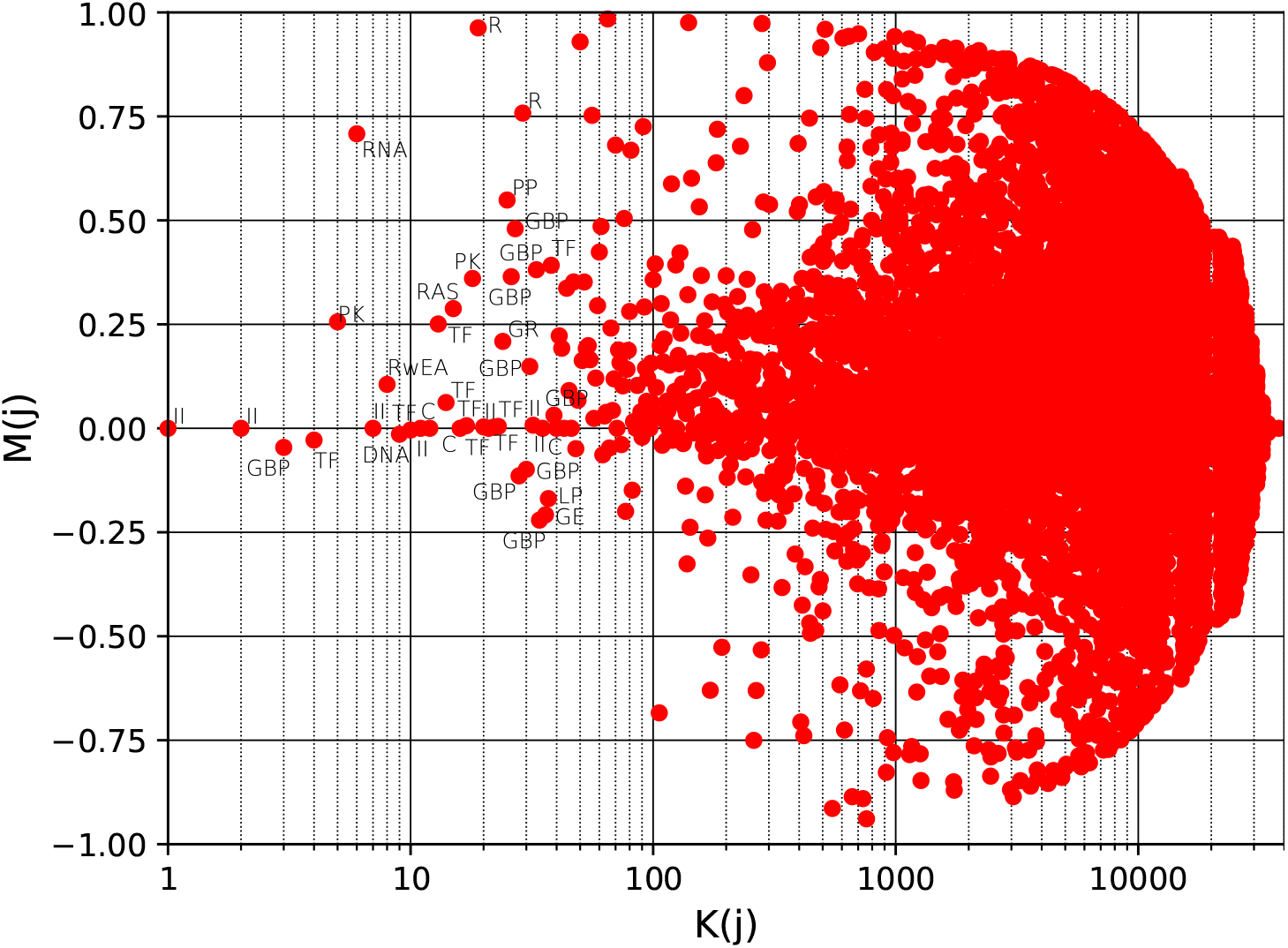
PageRank magnetization *M*(*j*) = (*P*_+_(*j*) – *P*− (*j*))/(*P*_+_(*j*) + *P* −(*j*)) for the Ising MetaCore network. Here, *j* is the node index and *K*(*j*) is the PageRank index of the node *j* in the simple Metacore network (without node doubling). The biological class is reported for the top 40 PageRank nodes (*K* ≤40, see Table 1): Inorganic ions (II), Generic binding protein (GBP), Transcription factor (TF), Protein kinase (PK), RNA, Receptor with enzyme activity (RwEA), DNA, Compound (C), RAS superfamily (RAS), Reaction (R), Generic receptor (GR), Protein phosphatase (PP), Generic enzyme (GE), Lipid phosphatase (LP).

Focusing on the top 40 PageRank in Fig. 3, we mainly observe that the nodes are either *non-magnetized M* ≈ 0, or positively magnetized *M* ≳ 1. These two situations correspond to biological elements which are equally activated/inhibited (*M* ≈ 0) and mainly activated (*M* ≳ 0), respectively. Among the top 40 PageRank nodes, the non-magnetized elements are mainly inorganic ions, such as H^+^, Na^+^, K^+^, Ca2^+^, and Cl^−^, which are involved in many elementary interactions. As non-magnetized nodes, we observe also very important biological molecules such as DNA and the ADP compound which should occupy a central place in the protein interaction network. Among positively magnetized nodes, we observe reactions (*M* ≳ 0.75), RNA (*M ≃* 0.7), protein kinase (*M ≃* 0.25 − 0.4) and phosphatase (*M ≃*0.55), which respectively are known to *turn on* and *turn off* proteins. Let us remark that, as DNA, RNA occupies a very central role in the protein interaction network (*K* = 6) but has a relatively high magnetization *M ≃*0.7 which indicates that RNA is mainly activated at the end of major biological processes. The other positively magnetized nodes correspond to some transcription factors, such as PPAR-gamma and STAT3, generic binding proteins, such as PI3K and GRB2, members of RAS superfamily, such as Rac1, and generic proteins, such as ITGB1. We nevertheless note that among the top 40 PageRank nodes, there are some mainly inhibited proteins (*M ≃*−0.2) such as the generic binding protein Bcl-2, the generic enzyme MDM2, and the lipid phosphatase PTEN.

We return to the magnetization properties of the selected subset of 12 nodes and the top 40 PageRank nodes in the next Section.

### 3.3 RIGMA analysis of the Ising MetaCore network

We illustrate the RIGMA analysis of the Ising MetaCore network by applying it to the subset of the 12 nodes given in Table 1. They are selected from the top 40 PageRank list of Table 1 by excluding simple molecules and keeping best ranked proteins according to PageRank probabilities. Each of the 12 nodes of the subset are doubled into a (+) component and a (−) component. We order these 24 nodes by ascending PageRank index *K* and alternating the (+) and the (−) components. This ordering is used to represent, in Fig. 4, the reduced Ising Google matrix *G*_R_ and its three matrix components *G*_rr_, *G*_pr_, and *G*_qr_. The weights of these components are respectively *W*_rr_ = 0.015, *W*_pr_ = 0.952, and *W*_qr_ = 0.033. As in the case of Wikipedia networks [12], the component *G*_pr_ has the highest weight, but as discussed, it is rather close to a matrix with identical columns, each one similar to the PageRank column vector. Thus, the *G*_pr_ matrix component does not provides more information than the standard PageRank/GMA analysis. We also see that the weight *W*_qr_ of the indirect links generated by long indirect pathways passing through the global Ising MetaCore network has approximately twice higher weight than the weight *W*_rr_ of direct links. Consequently, the contribution of indirect links are very important.

**Figure 4.**
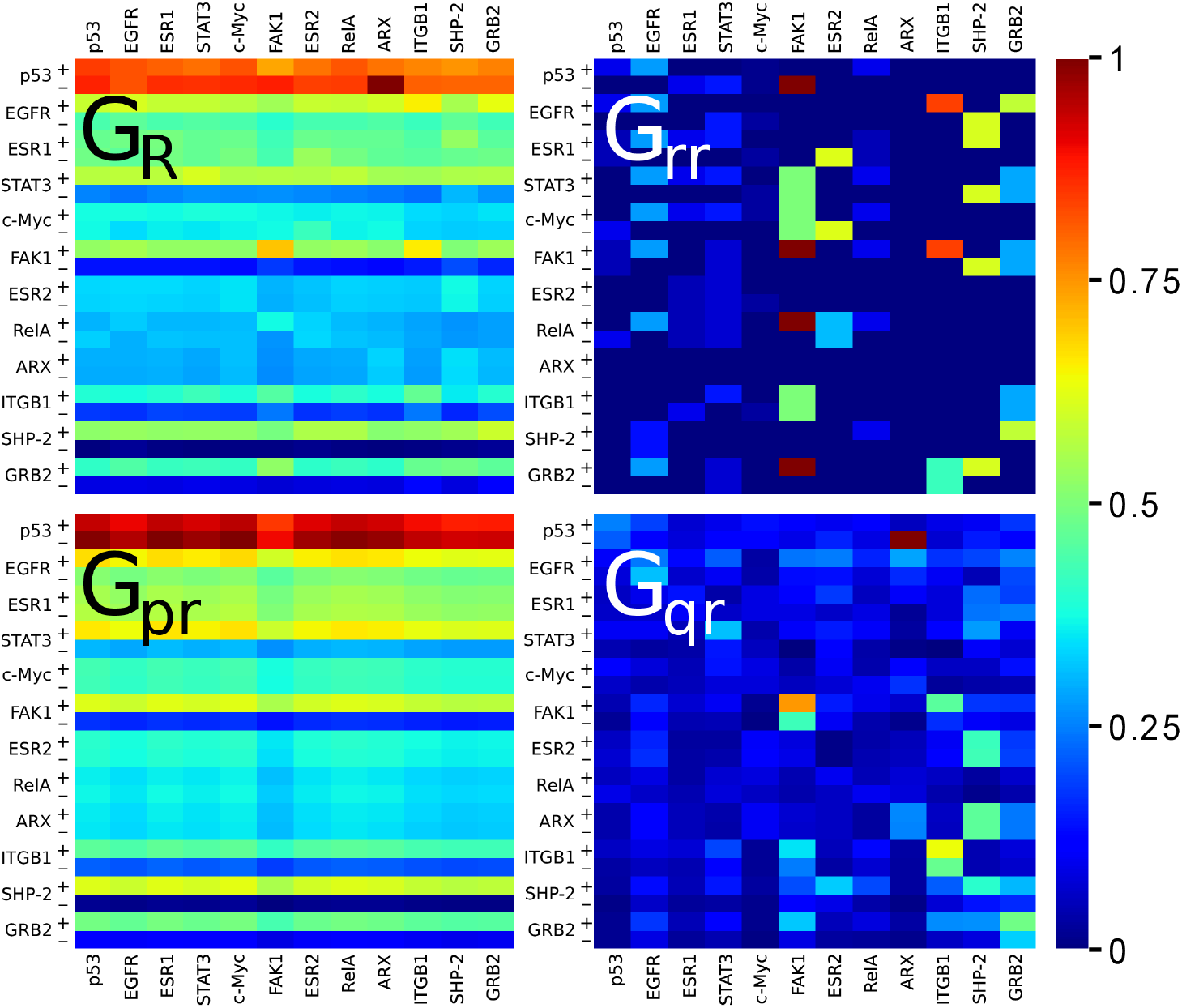
Reduced Google matrix *G*_R_ and its three matrix components *G*_pr_, *G*_rr_ and *G*_qr_ associated to the subset of nodes presented in Table 1 and belonging to the Ising MetaCore network. The weights of the matrix components are *W*_pr_ = 0.952, *W*_rr_ = 0.015, and *W*_qr_ = 0.033. Each colored cell corresponds to a *G*_X*ij*_ elements with X standing for R, rr, pr, or qr. A *G*_X*ij*_ matrix element is associated to the *j*→ *i* link where the *j* index corresponds to proteins read on the bottom or the top axis of the panels and the *i* index corresponds to proteins read on the left axis of the panels. The + and −signs correspond to the activated and inhibited state of the node, respectively. The values of the color bar correspond to the ratio of the matrix element over its maximum value. Note that the elements of *G*_qr_ may be possibly negative. There are only few and very small negative values (between –2.6× 10^−5^ and −5.3 ×10^−5^) which are not distinguishable from zero and have the same blue color code. Therefore, the color bar is only shown for positive values.

In the *G*_rr_ matrix component, each element *i* of the *j*th column corresponds to the direct action of the protein *j* on the protein *i*. The action is either an activation (+) or an inhibition (-). As a consequence, the *G*_rr_ matrix component simply mimics the Ising MetaCore network matrix adjacency (the elements of *G*_rr_ with a value equal to (greater than) (1 − *α*) /2*N* ≈ 0 correspond to values 0 (1 or 1/2) in the adjacency matrix of the Ising MetaCore network). It is interesting to compare the *G*_qr_ matrix elements with those of the *G*_rr_ matrix. Each one of the *G*_qr_ matrix elements either modifies, generally enhances, the weight of an existing link, for which a non zero matrix element exists in the *G*_rr_ matrix, or, interestingly, quantifies the strength of a hidden effective interaction between two proteins. As an example of the enhancement of an existing direct link, we observe, in Fig. 4, that the known activation of FAK1 by ITGB1 is enhanced by indirect links, ie, by pathways passing by the elements outside the set of the twelve chosen proteins. Also, we clearly observe also an enhancement of the self-activation of FAK1 and the appearance of its indirect self-inhibition.

Let us focus on the possible hidden interactions between the chosen set of twelve proteins. For that purpose, we show, in Fig. 5 (left panel), the matrix sum *G*_rr_ + *G*_qr_^(nd−block)^ which summarizes both the information concerning the direct and hidden interactions between the set of twelve proteins. Here, we use the *G*_qr_^(nd−block)^ matrix which is the *G*_qr_ matrix from which the diagonal elements (self-interaction terms) have been removed. In Fig. 5 (right panel), the *G*_rr_ + *G*_qr_^(nd−block)^ matrix elements associated to direct links have been masked to highlight only hidden interactions. Hence, although the ARX protein (aristaless related homeobox) is not directly connected to the other eleven proteins, ie, there is no direct action of the ARX protein onto the other eleven proteins and *vice versa*, it indirectly strongly inhibits the tumor suppressor protein p53. Secondarily, the ARX protein indirectly acts on different other proteins as it is indicated by blue shades on the ARX column in Fig. 5: hence, the ARX protein indirectly activates the EGFR and ESR1 proteins (epidermal growth factor receptor and estrogen receptor, respectively) and inhibits the c-Myc protein (proto-oncogene protein). Similarly, according to the *G*_rr_ matrix component (see Fig. 4), the c-Myc protein does not act on the chosen twelve proteins. But, the blues shades of the c-Myc column on the right panel of Fig. 5 gives us information on which proteins it indirectly contributes to activate or deactivate. Among strong weights of the *G*_rr_ + *G*_qr_^(nd−block)^ matrix sum, we observe also the SHP-2 phosphatase protein indirectly strongly interacts with with the ARX protein and the estrogen receptor protein ESR2. In return, the ESR2 protein, which directly inhibits ESR1 and c-Myc proteins, also indirectly activates the SHP-2 protein.

**Figure 5.**
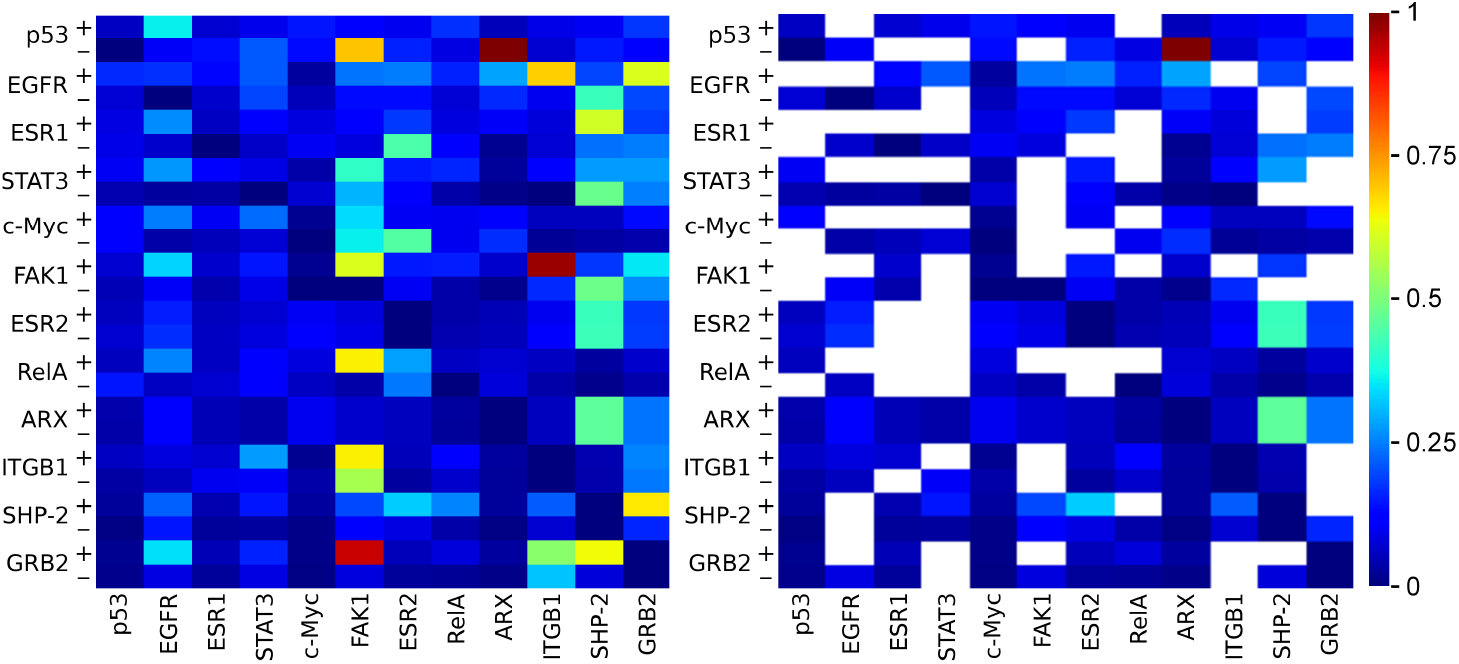
Sum of the two matrix components *G*_rr_ + *G*_qr_^(nd-block)^. The matrix components are the same as in Fig. 4 with the exception of *G*_qr_^(nd-block)^ which is obtained from *G*_qr_ by excluding 2 ×2 diagonal blocks, each one of these blocks corresponding to a protein self-loop. The right panel is the same as the left panel with the exception of the white cells which hide the direct links *j* →*i* between the 12 ×2 chosen nodes in the Ising MetaCore network. The values of the color bar correspond to the ratio of the matrix element over its maximum value.

In contrast to the adjacency matrix and the Google matrix, the matrix sum *G*_rr_+*G*_qr_ allows to discriminate the directed links outgoing from a given protein by assigning different weights to them. This discrimination is possible as the RGMA and the RIGMA takes account of not only the direct linkage of the twelve chosen proteins but all the knowledge encoded in the MetaCore complex network. Moreover, possible hidden links between proteins, which are non directly connected in the MetaCore network, can be inferred from non negligible weights in *G*_qr_. We propose to construct a reduced network highlighting the most important, direct and hidden, interactions between the twelve chosen proteins. Hence, for each protein *source* of the chosen subset, we retain, in the corresponding column of the *G*_rr_ + *G*_qr_ matrix, the two most important weights revealing the most important protein *target* of the protein *source*. Here, we do not consider self-inhibition and self-activation matrix elements in the matrix sum *G*_rr_ + *G*_qr_. The constructed reduced network associated to the twelve chosen proteins is presented in Fig. 6. We observe that it captures the above mentioned direct and hidden activation/inhibition actions between the considered proteins.

**Figure 6.**
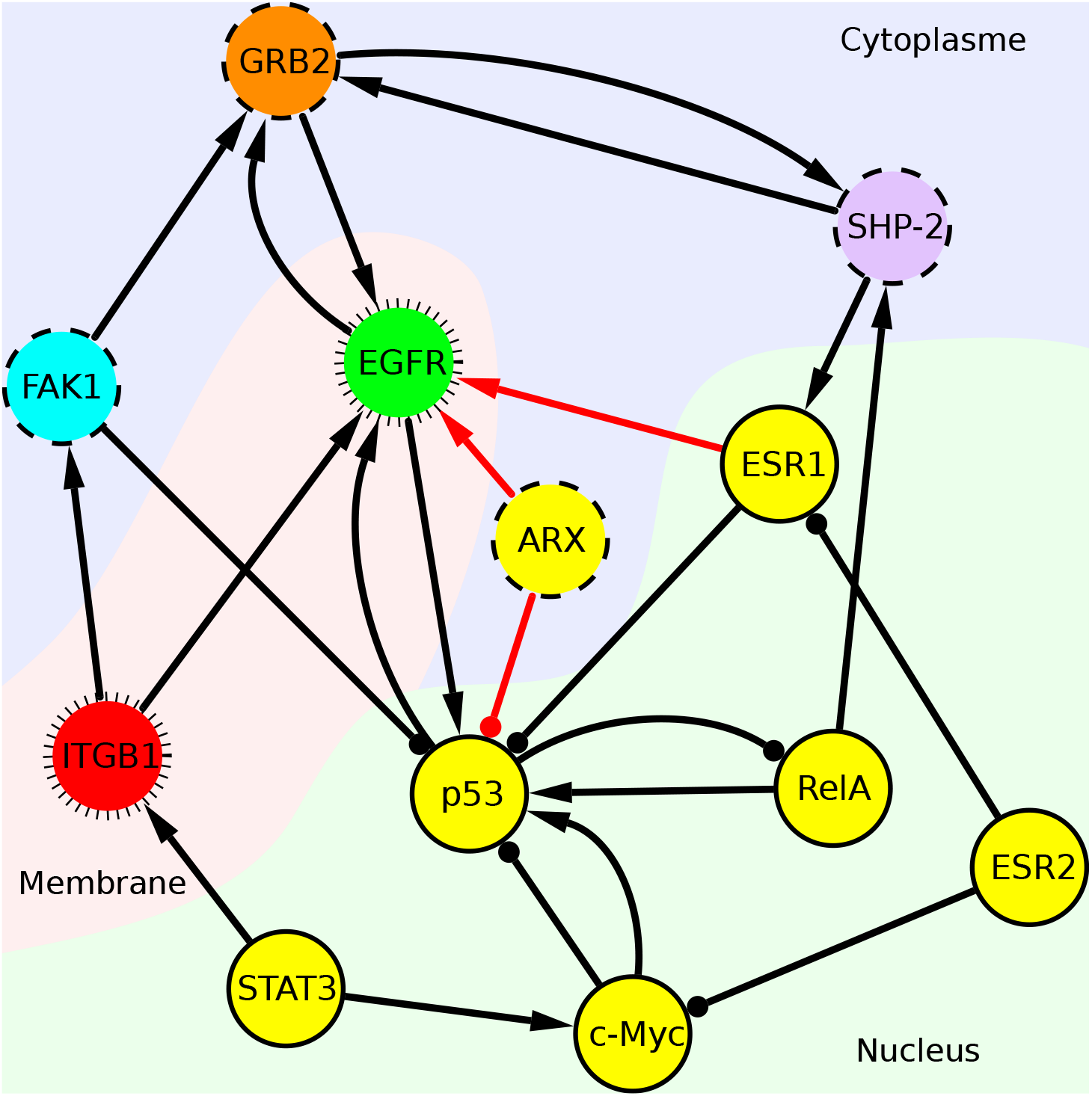
Reduced network of the chosen twelve proteins. (see Table 1). The construction procedure of this network is given in the main text. Arrow edges (→) represent activation links and dot edges (−•) represent inhibition links. The arrow tips and the dots are on the side of the target nodes. The black edges represent direct links. The red edges represent hidden links. The color of the nodes correspond to the type of proteins: transcription factors (yellow), protein kinase (cyan), generic receptor (red), receptor with enzyme activity (green), general binding protein (orange), and protein phosphatase (violet). The border style of the node correspond to the location of the proteins: nucleus (solid line), cytoplasm (dashed line), and plasma membrane (hairy line).

### 3.4 Sensitivity of the chosen subset

The PageRank sensitivity of the chosen subset of 12 proteins is obtained from the RIGMA and presented in Fig. 7 following the definitions given by (8) and (9). We remind that *F*_+_(*a, b*) gives the symmetric PageRank sensitivity of the nodes *a* and *b* to a variation of the link weight between them (in both directions from *a* to *b* and from *b* to *a*). The asymmetric PageRank sensitivity *F*_−_(*a, b*) determines what node is more sensitive to such weight variation. Thus, for *F*_−_(*a, b*) > 0 we obtain that node *a* is more influenced by node *b* and for *F*_−_(*a, b*) < 0 that node *b* is more influenced by node *a*.

**Figure 7.**
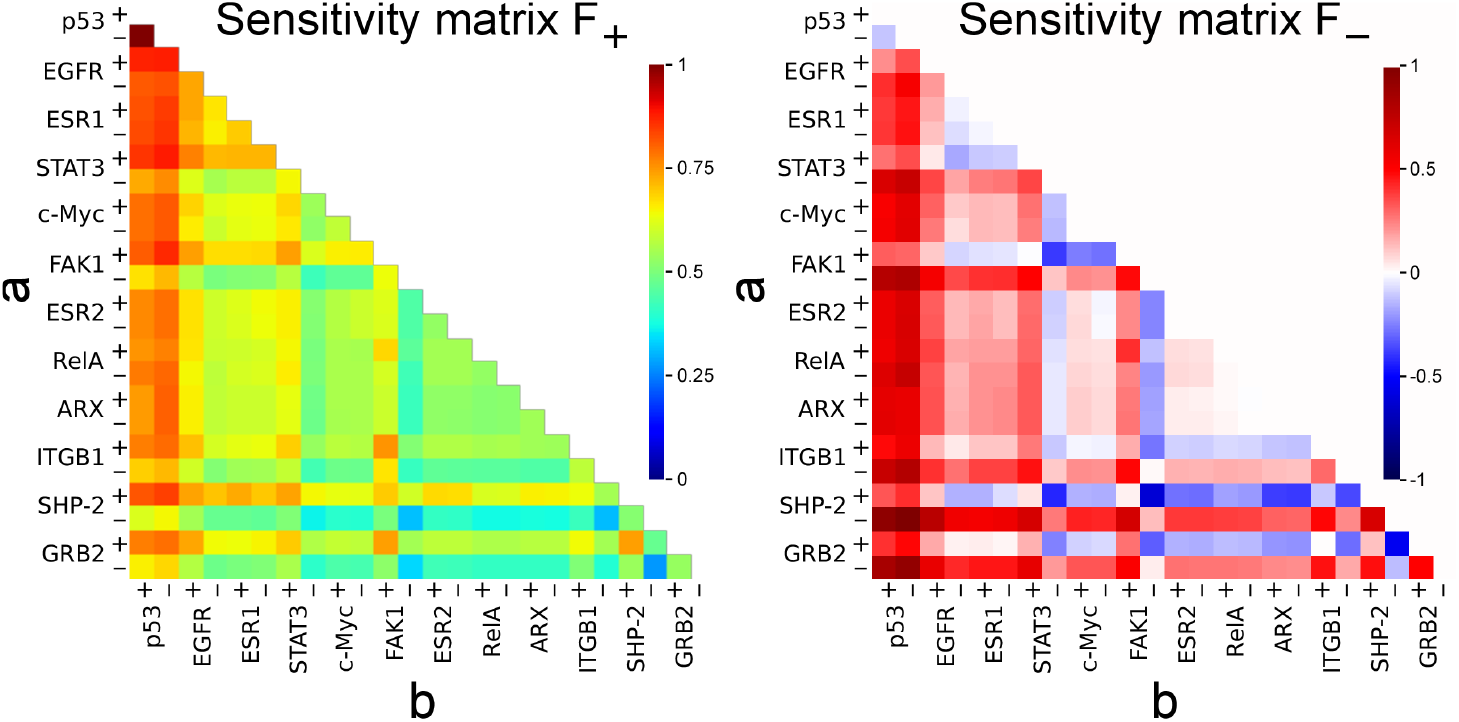
PageRank sensitivity matrices *F*_+_(*a, b*) (left panel) and *F* _−_ (*a, b*) (right panel) associated to the subset of nodes presented in Table 1 and belonging to the Ising MetaCore network. The values of the color bar correspond to *F*_+_(*a, b*)/ max_*a,b*_ (*F*_+_(*a, b*)) (left panel) and to *F*_−_(*a, b*)/ max_*a,b*_ |*F*_−_(*a, b*)| (right panel).

In Fig. 7, the symmetric PageRank sensitivity (left panel) shows that the activation or the inhibition of the p53 protein affect or are affected by all the other chosen proteins. Indeed, the p53 protein with *K* = 4 occupies a very central role in the protein interactions network as it contributes to the stability of the genome preventing damage biological information to be spread [22, 23, 24]. The reddish horizontal and vertical lines on the symmetric PageRank sensitivity panel (Fig. 7 left) indicate that the activation of the EGFR, STAT3, FAK1, SHP-2 and the GRB2 proteins are affected or affect all the other proteins of the chosen set. The right panel of the Fig. 7 shows the asymmetric PageRank sensitivity. We clearly observe that in fact it is the p53 protein which influences the activation/inhibition of the other proteins, and in a stronger manner the inhibition of the GRB2, SHP-2, ITGB1, and FAK1 proteins. In general, the inhibition of these four cited proteins is influenced by most of the other proteins (see greenish horizontal lines) and in return their respective activation influences also the other proteins (see greenish vertical lines).

### 3.5 Examples of magnetization of nodes

In Fig. 8, left panel, we show, in the PageRank-CheiRank (*k, k*^*^)-plane (see Tables 1 and 2 for the relative PageRank and CheiRank indexes *k* and *k*^*^), the PageRank magnetization *M* of the chosen 12 proteins. These nodes have global PageRank indexes *K* ≤ 26 (see Table 1). In agreement with data presented in Fig. 3, for such *K* values, the magnetization is indeed mainly positive. So, these proteins are primarily activated. More precisely, as they belong to the top PageRank of the proteome (*K*/*N* ≤ 0.6%_0_), these proteins are activated as the result of most important cascade of interactions along the causality pathways. The magnetization *M* is also presented in Fig. 7, right panel, but for every nodes with *K* ≤ 40 (see Table 1). Here, for these top PageRank indexes, we have both positive and negative magnetization values, but the majority of the nodes have a magnetization close to zero, as discussed in Fig. 3.

**Figure 8.**
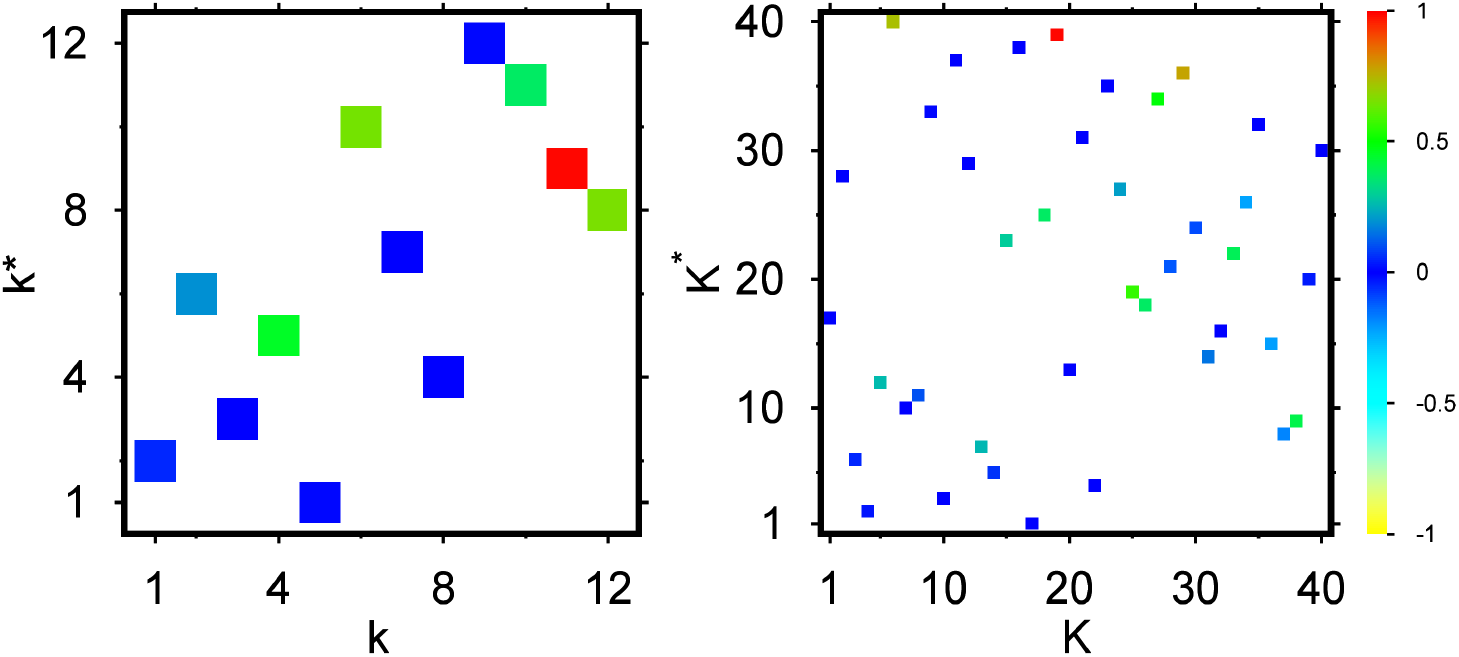
PageRank magnetization *M*(*K*) = (*P*_+_(*K*) − *P*_–_ (*K*))/(*P*_+_(*K*) + *P*_−_ (*K*)) presented in the PageRank-CheiRank (*K, K*^*^)-plane. Left panel: PageRank magnetization *M*(*k*) for the chosen twelve proteins presented in the relative indexes (*k, k*^*^)-plane (see *k* adn *k*^*^ indexes in Table 1). Right panel: PageRank magnetization *M*(*K*) for nodes with *K* ≤ 40. Here, *P*_±_(*K*) is the PageRank probability of the (±) component of the Ising MetaCore network node associated with the *K* PageRank (see text). The values of the color bar correspond to *M*/ max |*M*| with max_*k*≤12_ |*M*(*k*)| = 0.549 (left panel) and max_*K*≤40_ |*M*(*K*)| = 0.963 (right panel). On the right panel, the *K*^*^ index is here the relative CheiRank index inside the set of the first *K* ≤ 40 nodes.

For the top 40 PageRank (*K* ≤ 40), the top 3 most activated nodes are the K^+^ Potassium ion in cytosol (*K* = 19, *M*(*K*) = 0.962815), the 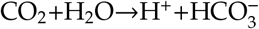 reaction (*K* = 29, *M*(*K*) = 0.757892), and the intracellular mRNA (*K* = 6, *M*(*K*) = 0.708154), and the top 3 most inhibited nodes are the generic binding protein Bcl-2 (*K* = 34, *M*(*K*) = −0.220751), the generic enzyme MDM2 (*K* = 36, *M*(*K*) = −0.208082), and the generic binding protein E-cadherin (*K* = 28, *M*(*K*) = −0.114311).

## 4 Discussion

In this work, we have presented a detailed description of the statistical properties of the protein-protein interactions MetaCore network obtained with extensive Google matrix analysis. In this way, we find the proteins and molecules which are at the top PageRank and CheiRank positions playing thus an important role in the influence flow through the whole network structure. With a simple example of a subset of selected proteins (subset of selected nodes), we show that the reduced Google matrix analysis (RGMA) allows to determine the effective interactions between these proteins taking into account all the indirect pathways between these proteins through the global MetaCore network, in addition to direct interactions between selected proteins. We stress that the approach with the reduced Ising Google matrix algorithm, based on Ising spin description, allows to take into account the bifunctional nature of the protein-protein interactions (activation or inhibition) and to determine the average action type (or magnetization) of each protein.

Here, we have presented mainly the statistical properties of the MetaCore network without entering into detailed analysis of related biological effects. We plan to address, in further studies, the biological effects obtained from the reduced Google matrix analysis of the MetaCore network.

## Abbreviation

PPI: protein-protein interactions
GMA: Google matrix analysis
IGMA: Ising Google matrix analysis
RGMA: reduced Google matrix analysis
RIGMA: reduced Ising Google matrix analysis
WWW: World Wide Web

## Competing interests

The authors declare that they have no competing interests.

## Author’s contributions

The authors contributed equally to this work. All authors read and approved the final manuscript.

## Acknowledgements

We thank Andrei Zinovyev (Institut Curie) for useful discussions.

## Funding

This research has been partially supported through the grant NANOX N° ANR-17-EURE-0009 (project MTDINA) in the frame of the Programme Investissements d’Avenir, France. This research has been also supported by the Programme Investissements d’Avenir ANR-15-IDEX-0003, ISITE-BFC (GNETWORKS project) and the council of Bourgogne Franche-Comté region (APEX project and REpTILs project).

## Notes

http://quantware.ups-tlse.fr/QWLIB/metacorenet/

### Competing Interest Statement

The authors have declared no competing interest.

